# Atomistic Insights into TRPC6–Caveolin-1 Interactions Interface via All-Atom Molecular Dynamics Simulations: Structural and Energetic Basis for Selective Modulation

**DOI:** 10.64898/2026.09.09.750526

**Authors:** Shayesteh Bazsefidpar, Alejandro Rodriguez, Masayuki X Mori, Hamid Mosadeghi, Ajay K Israni, Kyle M Koss, Onur K Polat

## Abstract

The molecular determinants governing TRPC6 stabilization and its interaction with caveolin-1 (CAV1) remain poorly defined, despite their critical role in caveolae organization and signaling. The absence of atomistic structural models has hindered a mechanistic understanding of how TRPC6 is recruited to, and stabilized within, caveolar microdomains. Here, we combine all-atom molecular dynamics simulations with MM/PBSA calculations to characterize the TRPC6–CAV1 complex at atomic resolution. To preserve a biologically realistic membrane environment, harmonic restraints were applied to transmembrane and membrane-embedded regions of both proteins, while the cytosolic TRPC6 N-terminus and solvent-exposed edges of the CAV1 scaffolding domain were kept fully flexible, allowing the putative caveolin-binding motif to explore conformational space and form dynamic contacts. This protocol maintained overall structural integrity while capturing physiologically relevant flexibility at the interaction surface. MM/PBSA analysis revealed a highly favorable binding free energy (ΔG_binding = −255.9 ± 1.6 kJ/mol), dominated by electrostatic contributions and reinforced by hydrophobic and aromatic interactions. Per-residue energy decomposition identified an acidic patch in TRPC6 (residues 30–41) that engages a complementary basic, amphipathic segment in CAV1 (residues 85–106), defining a cooperative, reversible binding interface. These findings provide the first atomistic description of TRPC6 recruitment by CAV1 and establish a quantitative framework for the rational design of strategies to selectively modulate this interaction.

**Highlight:**

- Established the first atomistic model of the TRPC6–CAV1 interaction in caveolae
- Combined all-atom MD simulations with restrained membrane-embedded protein models
- Interface analysis shows reversible noncovalent binding via transient H-bonds and salt bridges
- MM/PBSA revealed strong electrostatic-driven binding of the TRPC6–CAV1 complex
- Identified an acidic TRPC6 N-terminal hotspot engaging a basic amphipathic CAV1 region

**Graphical Abstract:** 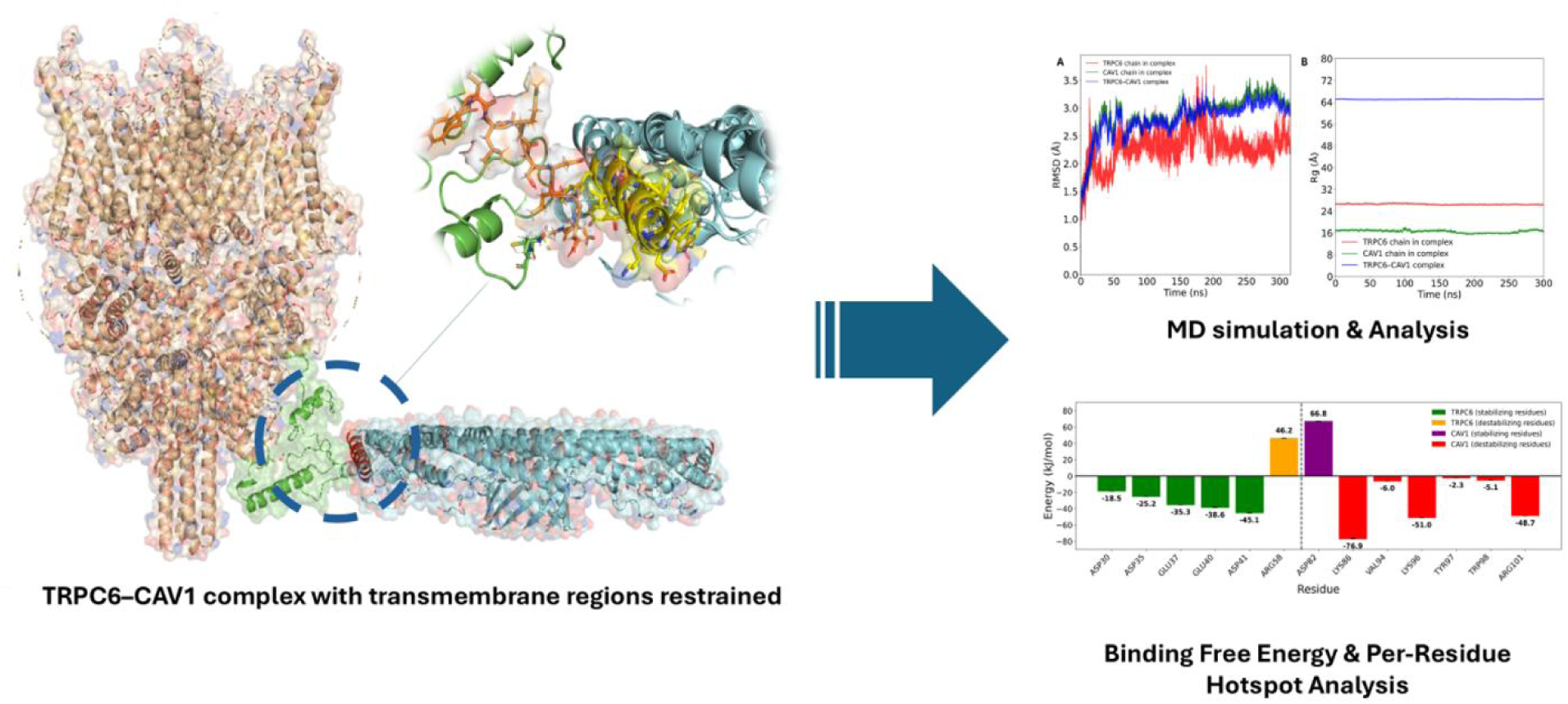

## 1. Introduction

Calcium signaling fundamentally regulates cellular homeostasis, governing mechanotransduction, cytoskeletal dynamics, and vascular function [1]r normal physiology, while sustained or mislocalized calcium influx triggers pathological cascades [2]. Membrane microdomains—particularly caveolae—have emerged as critical organizers of calcium signaling by concentrating ion channels and effectors into specialized functional platforms[1].

The Transient Receptor Potential Canonical 6 (TRPC6) channel is a non-selective cation channel that mediates calcium entry in response to G protein–coupled receptor activation and mechanical stimuli [3]. TRPC6 plays a pivotal role in mechanosensitive signaling in vascular cells and kidney podocytes [4], where precise control of channel localization and activity is required to maintain cytoskeletal integrity and filtration function [5]. Aberrant TRPC6 activity has been directly linked to human kidney diseases, including focal segmental glomerulosclerosis, in which sustained calcium influx promotes actin cytoskeleton remodeling, podocyte injury, and proteinuria [6]. These observations highlight the importance of regulatory mechanisms that govern TRPC6 spatial organization at the plasma membrane.

Caveolae are cholesterol- and sphingolipid-enriched, flask-shaped plasma membrane invaginations that serve as mechanosensory and signaling platforms [7]. Caveolin-1 (CAV1), their principal structural protein, also functions as a scaffolding adaptor that recruits and organizes diverse signaling effectors [8,9]. Through its caveolin scaffolding domain (CSD), CAV1 engages caveolin-binding motifs (CBMs) in target proteins, enabling dynamic, reversible protein–protein interactions at the cytoplasmic face of caveolar membranes [10]. These interactions precisely regulate the localization, stability, and activity of multiple signaling proteins, including ion channels [2]. Importantly, mutations in CAV1 and dysregulation of caveolae contribute to the development and progression of diseases such as cancer, asthma, pulmonary fibrosis, pulmonary arterial hypertension, chronic inflammatory respiratory diseases, and lipodystrophy. Mutations such as P132L interfere with CAV1 oligomerization and caveolae assembly, highlighting the structural and functional importance of CAV1 in health and disease [11].

Although several TRPC family members—most notably TRPC1—have been experimentally shown to associate with CAV1 [12–14]. Moreover, studies on TRPC5 have shown that caveolin-1 acts as a dynamic scaffolding protein, assembling TRPC channels with signaling partners such as eNOS to coordinate Ca²⁺ influx and nitric oxide signaling. While disruption of caveolin-binding domains in TRPC5 markedly impairs its association with CAV1 and downstream signaling, the structural determinants underlying this regulation remain unresolved, underscoring the need for atomistic insight into caveolin-dependent TRPC channel regulations [15]. However, these studies provide limited structural insight into the molecular determinants governing caveolin–TRPC recognition. In this context, the structural basis of the TRPC6–CAV1 interaction remains unresolved [16]. Available cryo-electron microscopy structures of TRPC6 lack the flexible cytoplasmic regions predicted to harbor caveolin-binding motifs, precluding direct experimental visualization of the interaction interface [16]. Consequently, the molecular determinants governing TRPC6 recruitment to caveolae and its regulation by CAV1 remain poorly understood [17]. Emerging evidence implicates excessive TRPC6 stabilization within caveolar microdomains as a driver of pathological calcium accumulation in human kidney cells through prolonged channel activity [18]. Direct pharmacological blockade of TRPC6 ion conduction remains challenging due to the channel’s essential physiological roles and potential for off-target effects across the TRP family [15,19]. Furthermore, loss of function of TRPC6 channels also lead to kidney injury [20]).In contrast, selectively disrupting protein–protein interactions that govern TRPC6 localization and signaling offers a precise therapeutic strategy [21] Targeting the TRPC6–CAV1 interaction thus holds promise for attenuating disease-associated calcium influx while preserving baseline channel function. Targeting the TRPC6–CAV1 interaction thus holds promise for attenuating disease-associated calcium influx while preserving baseline channel function.

Peptide-based inhibitors targeting caveolin scaffolding domain recognition have emerged as promising modulators of caveolae-dependent signaling pathways [22,23]. However, their rational design demands detailed structural characterization of the TRPC6–CAV1 interaction interface and identification of the key molecular determinants driving complex formation and stability.

To address this critical knowledge gap, we employed an integrative computational approach that combines artificial intelligence–based structure prediction with molecular dynamics simulations to explore the conformational landscape and interaction mechanism of the TRPC6–CAV1 complex. Binding free energies were quantified and complemented by per-residue energy decomposition analyses to identify key energetic contributors to complex stabilization. By resolving electrostatic and aromatic interaction hotspots at the interface, this study provides an atomistic and energetic framework for the rational design of therapeutic peptides targeting disruption of the TRPC6–CAV1 interaction.

## 2. Materials and methods

### 2.1. N-terminal Modeling of TRPC6

The N-terminal region of TRPC6 (residues 1–84), absent from available experimental structures, was modeled using ESMFold, an AI-driven structure prediction method that leverages language-model embeddings from ESM-2 to generate atomic-level coordinates directly from sequence [24]. The amino acid sequence was obtained from UniProt entry Q9Y210 (TRPC6_HUMAN). The predicted model was refined with secondary structure assignments and used as the starting point for downstream molecular dynamics simulations.

### 2.2. Molecular dynamics simulation setup

All simulations were performed using were performed using GROMACS 2024.2 with PLUMED 2.9.2, executed on an HPC cluster equipped with NVIDIA A100 GPUs. The crystal structure of TRPC6 [PDB: 6UZA] and Caveolin-1 [PDB: 7SC0] was obtained from the RCSB Protein Data Bank. The system was solvated in a cubic water box, and chloride (Cl⁻) and sodium (Na⁺) ions were added to neutralize the total charge. Atomic interactions were modeled using the CHARMM36. The TIP3P water model [25] was selected for solvation. Energy minimization was carried out using the steepest-descent algorithm (up to 5000 steps, convergence criterion 1000 kJ/mol/nm) with a Verlet cutoff scheme, 1.2 nm real-space cutoffs for both electrostatic and van der Waals interactions, and Particle Mesh Ewald (PME) for long-range electrostatics. Equilibration proceeded in two phases: a 1 ns NVT run at 300 K using the V-rescale thermostat (coupling time 0.1 ps), followed by a 2 ns NPT run at 1 bar using the C-rescale barostat (coupling time 2.0 ps). Long-range electrostatics were treated with PME throughout, and all covalent bonds were constrained using LINCS to allow a stable integration timestep. Trajectory analysis was performed with GROMACS utilities. Structural stability was assessed from the backbone rootmeansquare deviation (RMSD), and complex compactness was quantified using the radius of gyration (Rg). In addition, the time evolution of intermolecular hydrogen bonds between TRPC6 and CAV1 was quantified to evaluate interface Energy minimization was carried out using the steepest-descent algorithm (up to 5000 steps, convergence criterion 1000 kJ/mol/nm) with a Verlet cutoff scheme, 1.2 nm real-space cutoffs for both electrostatic and van der Waals interactions, and Particle Mesh Ewald (PME) for long-range electrostatics. Equilibration proceeded in two phases: a 1 ns NVT run at 300 K using the V-rescale thermostat (coupling time 0.1 ps), followed by a 2 ns NPT run at 1 bar using the C-rescale barostat (coupling time 2.0 ps). Long-range electrostatics were treated with PME throughout, and all covalent bonds were constrained using LINCS to allow a stable integration timestep. Trajectory analysis was performed with GROMACS utilities. Structural stability was assessed from the backbone rootmeansquare deviation (RMSD), and complex compactness was quantified using the radius of gyration (Rg). In addition, the time evolution of intermolecular hydrogen bonds between TRPC6 and CAV1 was quantified to evaluate interface Energy minimization was carried out using the steepest-descent algorithm (up to 5000 steps, convergence criterion 1000 kJ/mol/nm) with a Verlet cutoff scheme, 1.2 nm real-space cutoffs for both electrostatic and van der Waals interactions, and Particle Mesh Ewald (PME) for long-range electrostatics. Equilibration proceeded in two phases: a 1 ns NVT run at 300 K using the V-rescale thermostat (coupling time 0.1 ps), followed by a 2 ns NPT run at 1 bar using the C-rescale barostat (coupling time 2.0 ps). Long-range electrostatics were treated with PME throughout, and all covalent bonds were constrained using LINCS to allow a stable integration timestep. Trajectory analysis was performed with GROMACS utilities. Structural stability was assessed from the backbone rootmeansquare deviation (RMSD), and complex compactness was quantified using the radius of gyration (Rg). In addition, the time evolution of intermolecular hydrogen bonds between TRPC6 and CAV1 was quantified to evaluate interface Energy minimization was carried out using the steepest-descent algorithm (up to 5000 steps, convergence criterion 1000 kJ/mol/nm) with a Verlet cutoff scheme, 1.2 nm real-space cutoffs for both electrostatic and van der Waals interactions, and Particle Mesh Ewald (PME) for long-range electrostatics. Equilibration proceeded in two phases: a 1 ns NVT run at 300 K using the V-rescale thermostat (coupling time 0.1 ps), followed by a 2 ns NPT run at 1 bar using the C-rescale barostat (coupling time 2.0 ps). Long-range electrostatics were treated with PME throughout, and all covalent bonds were constrained using LINCS to allow a stable integration timestep. Trajectory analysis was performed with GROMACS utilities. Structural stability was assessed from the backbone root-mean-square deviation (RMSD), and complex compactness was quantified using the radius of gyration (Rg). In addition, the time evolution of intermolecular hydrogen bonds between TRPC6 and CAV1 was quantified to evaluate interface stability.- mean-square deviation (RMSD), and complex compactness was quantified using the radius of gyration (Rg). Solvent-accessible surface area (SASA) was computed to monitor changes in solvent exposure and detect conformational rearrangements. In addition, the time evolution of intermolecular hydrogen bonds between TRPC6 and CAV1 was quantified to evaluate interface stability.

### 2.3. Binding Free Energy Estimation Using MM-PBSA

The Molecular Mechanics Poisson–Boltzmann Surface Area (MM/PBSA) approach is a widely used method in computational drug discovery, providing detailed insights into the energetic contributions of protein–ligand interactions [10]. This method decomposes binding free energy into several components, including van der Waals interactions, electrostatic interactions, polar solvation energy, and non-polar solvation energy estimated from solvent-accessible surface area (SASA). Consequently, MM/PBSA enables identification of the key forces driving binding affinity [26]. In this study, MM/PBSA analysis was performed using the g_mmpbsa package (https://rashmikumari.github.io/g_mmpbsa/). Calculations were conducted using a single-step approach based on protein–protein molecular dynamics trajectories and their corresponding parameter files. The MmPbsaStat.py script included in the g_mmpbsa package was employed to compute individual energy components, sampling configurations every 100 ps from 50 ns to the end of each MD trajectory. The total binding free energy (ΔG_binding) was calculated as the sum of the van der Waals (ΔE_vdW), electrostatic (ΔE_elec), polar solvation (ΔG_polar), and non-polar solvation (ΔG_nonpolar) energy contributions. Furthermore, to identify key residues contributing to binding, we calculated per-residue energy decomposition using the same g_mmpbsa workflow. This analysis allowed us to determine which amino acid residues in the binding site contribute most significantly to the binding affinity, thereby providing mechanistic insights into the interaction.

## 3. Results and Discussion

### 3.1. N-terminal Modeling of TRPC6 and Pre-equilibration

The N-terminal region of TRPC6 (residues 1–84), which is absent from available cryo-EM structures, was modeled using ESMFold, yielding a high-confidence structure with pLDDT scores exceeding 85 for more than 95% of residues (Fig. 1). Secondary structure prediction using PSIPRED identified two α-helices (H1: residues 17–27; H2: residues 61–86) separated by flexible loop regions, in agreement with the ESMFold model.

**Fig. 1.**
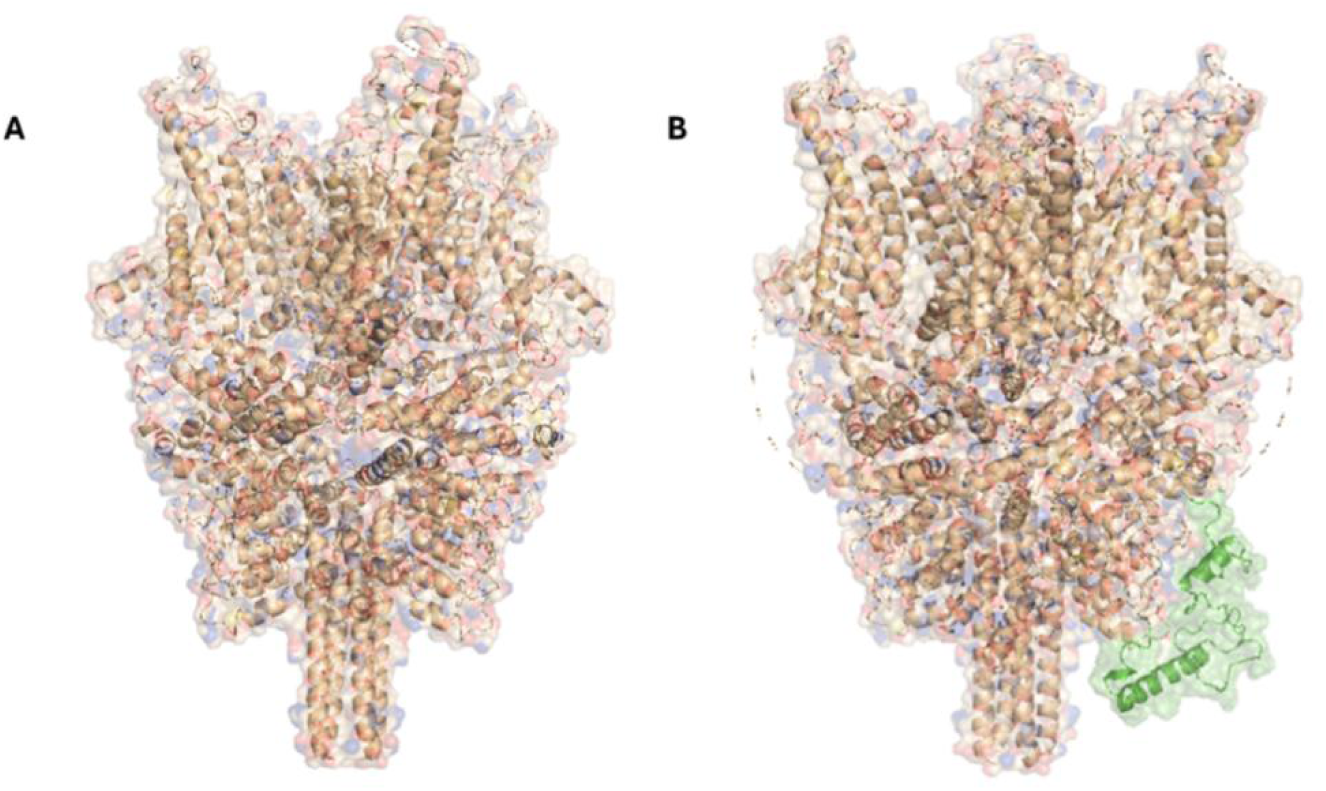
ESMFold modeling of the TRPC6 N-terminus. (A) N-terminal segment before modeling. (B) Final model with defined secondary structure

To assess structural stability, the modeled N-terminal domain was subjected to 500 ns of all-atom MD simulation in explicit solvent using the CHARMM36m force field. Throughout the simulation, the backbone RMSD fluctuated between 7 and 11 Å, reflecting the intrinsic flexibility of this isolated N-terminal region and the reorientation of secondary structural elements rather than global unfolding. Importantly, the radius of gyration (Rg) rapidly stabilized at approximately 21 Å and remained constant over the course of the simulation, indicating maintenance of a compact and well-packed fold (Fig. 2A,B).

**Fig. 2.**
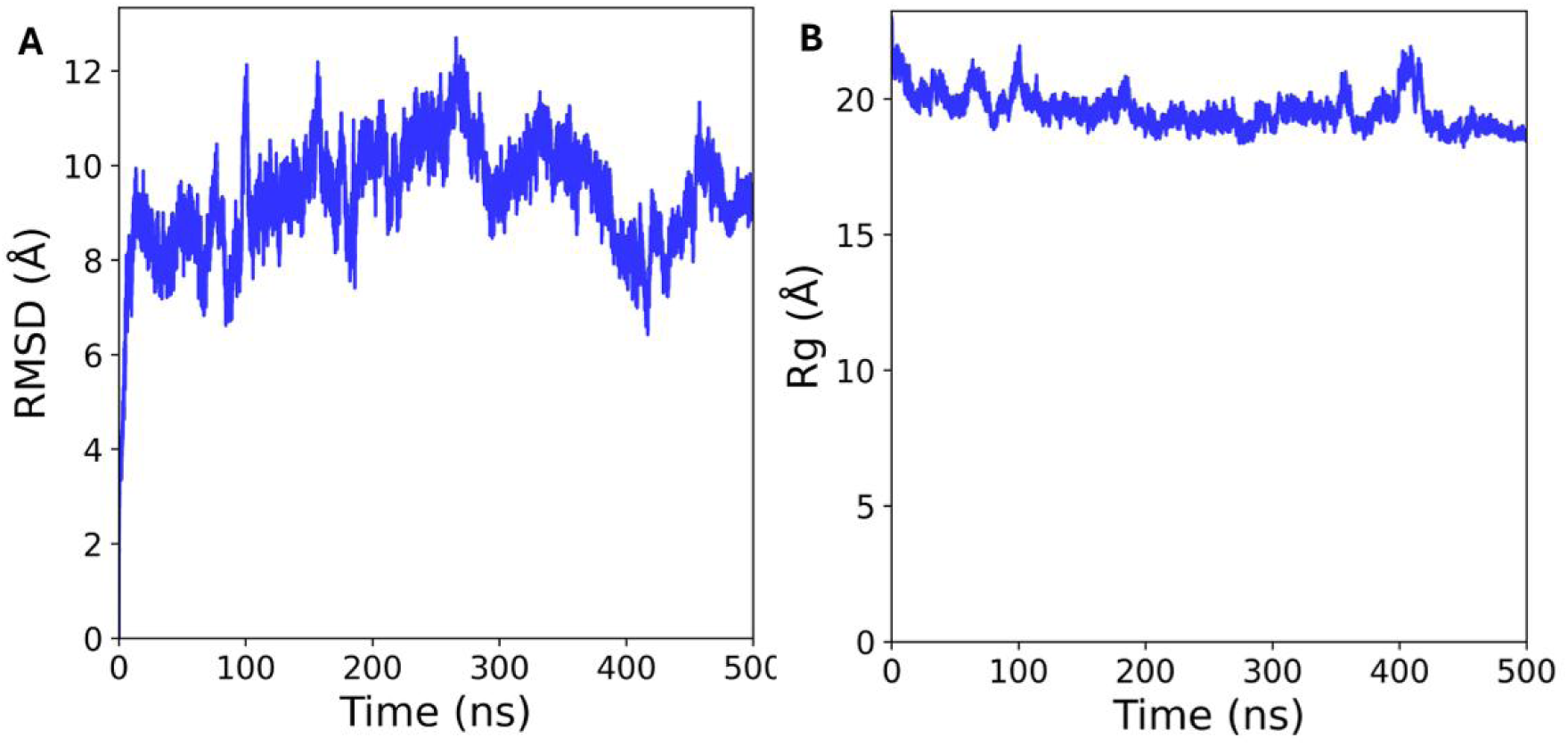
Structural stability of the TRPC6 N-terminal domain during MD simulation, assessed by (A) backbone RMSD and (B) Rg, confirming equilibration prior to complex assembly.

Together, these results demonstrate that despite its dynamic nature, the TRPC6 N-terminal domain retains structural integrity and behaves as a stable scaffold suitable for protein– protein interactions. The equilibrated conformation obtained from MD simulations therefore provides a reliable structural foundation for subsequent docking with the CAV1 region and for further simulations aimed at elucidating the molecular basis of TRPC6–CAV1 recognition.

### 3.2. Molecular Dynamics Simulation of TRCP6-CAV1 Complex

The caveolin signaling hypothesis posits that caveolins serve as dynamic scaffolding proteins that recruit and organize diverse signaling molecules within caveolar membrane microdomains [27,28]. Caveolae comprise polyhedral membrane invaginations stabilized by a flexible cavin-1 network surrounding oligomeric CAV1 discs. Cryo-EM reconstructions of the human CAV1 8S complex reveal a spiral assembly of 11 protomers forming a compact ∼15 nm disc [29–31]. One disc surface remains relatively flat and hydrophobic—consistent with membrane association—while the opposing face features a raised rim (∼3 nm deep, spanning two helices) and a central protruding β-barrel, with the outer rim likewise exhibiting hydrophobic character to facilitate lipid interactions [31,33]. Early biochemical studies identified the CAV1 scaffolding domain (CSD; residues 81–101) within this outer rim as a critical interaction module that recognizes caveolin-binding motifs (CBMs), typically enriched in aromatic residues. These interactions are characteristically transient, non-covalent, and reversible—properties essential for regulated cellular signaling [27,33].

**Fig. 3** shows an overall view of the TRPC6–CAV1 complex used in the simulations. To maintain physiological membrane orientation during MD simulations, CAV1 was embedded in a membrane-associated configuration with harmonic positional restraints applied to transmembrane and membrane-embedded residues. These restraints stabilized protein orientation relative to the lipid bilayer while preventing unphysical displacement or rotation. Critically, residues at the cytosolic interface of the caveolin scaffolding domain (CSD) were left unrestrained to preserve functionally relevant flexibility for protein–protein interactions. Also, a comparable restraint scheme was implemented for TRPC6. Residues within or adjacent to transmembrane segments were restrained to maintain channel topology with respect to the lipid bilayer, whereas the cytosolic Also, a comparable restraint scheme was implemented for TRPC6. Residues within or adjacent to transmembrane segments were restrained to maintain channel topology with respect to the lipid bilayer, whereas the cytosolic N-terminal region—previously implicated in caveolin recognition [13,34], was left unrestrained. This approach enabled the caveolin-binding motif (CBM)-containing segment of TRPC6 to sample conformational space freely and engage the CAV1 CSD during the simulation, thereby capturing physiologically relevant interaction dynamics.

**Fig 3.**
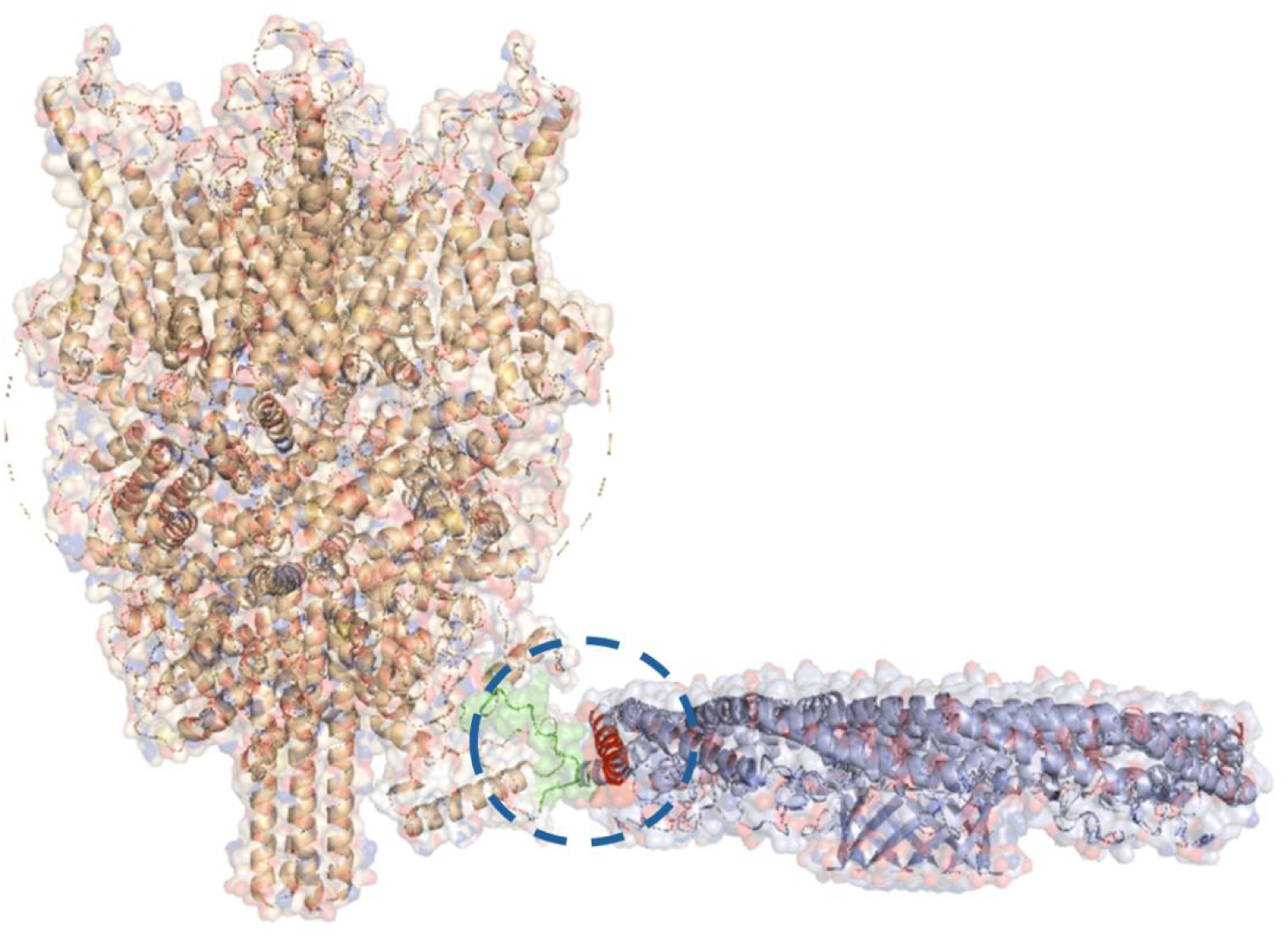
Overview of the TRPC6–CAV1 complex used for molecular dynamics simulations, showing TRPC6 positioned relative to the oligomeric CAV1 disc. Transmembrane and membrane-embedded regions of both TRPC6 and CAV1 were subjected to harmonic positional restraints to preserve physiological membrane orientation, whereas the cytosolic TRPC6 N-terminus and the cytosolic face of the CAV1 scaffolding domain were left unrestrained to allow dynamic protein–protein interactions.

### 3.3. Interface and binding analysis of protein-protein complex

To elucidate the structural determinants underlying the TRPC6–CAV1 interaction, interface and binding energy parameters were analyzed using PDBePISA (https://www.ebi.ac.uk/pdbe/pisa/). The analysis encompassed atomic and residue composition at the interface, solvent-accessible surface area (SASA), solvation energy changes upon complex formation, and statistical significance (P-values) (Table 1). The TRPC6–CAV1 interface exhibited geometric complementarity, comprising 92 atoms from TRPC6 (9.1% of surface atoms) and 82 atoms from CAV1 (9.1% of surface atoms), corresponding to 27 residues (5.3% of total TRPC6 residues) and 18 residues (16.1% of total CAV1 residues), respectively. This interface made substantial contributions to complex stabilization, burying solvent-accessible surface areas of 749.0 Å² for TRPC6 (∼2.8% of total SASA) and 796.1 Å² for CAV1 (∼10.5% of total SASA). Solvation energy analysis revealed favorable gains upon complex formation (−2.1 kcal/mol for TRPC6; −5.3 kcal/mol for CAV1), with average per-residue contributions of −2.7 kcal/mol (TRPC6) and −2.2 kcal/mol (CAV1)—values consistent with a stable, energetically favorable association. The Complexation Significance Score (CSS) ranged from 0.312 to 0.616, aligning with biologically relevant protein–protein interfaces. Statistical validation yielded P-values of 0.146 (TRPC6) and 0.353 (CAV1), confirming that the observed inter-protein contacts significantly exceed expectations from random molecular collisions. Collectively, these quantitative metrics establish the TRPC6–CAV1 interface as a functionally relevant, energetically stable protein–protein interaction characterized by extensive buried surface area, favorable desolvation energetics, and statistically significant residue-level contacts.

**Table 1.** Analysis of TRPC6–CAV1 Surface Interactions: Structural and Energetic Properties.

| Property | TRPC6–CAV1 |
| --- | --- |
| Class | Protein-Protein |
| Number of atoms (Interface) | 92 (9.1 %) - 82 (9.1 %) |
| Number of atoms (Surface) | 2412 (62.8 %) – 606 (67.1 %) |
| Number of atoms (Total) | 3839 - 903 |
| Number of residues (Interface) | 27 (5.3 %) – 18 (16.1 %) |
| Number of residues (Surface) | 475 (93.3 %) - 106 (94.6 %) |
| Number of residues (Total) | 509 - 112 |
| Solvent-accessible area (Interface) (Å<sup>2</sup>) | 749.0 – 796.1 |
| Solvent-accessible area (Total) (Å<sup>2</sup>) | 26935.7 - 7561.9 |
| Solvation energy (Isolated Structure) (kcal/mol) | −415.9 - −71.5 |
| Solvation energy (Gain on Complex Formation)<br>(kcal/mol) | -2.1 (0.5 %) - -5.3 (7.5 %) |
| Solvation energy (Average Gain) (kcal/mol) | -2.7 (3.6 %) - -2.2 (16.6 %) |
| CSS (Complexation Significance Score) | 0.616 - 0.312 |
| P-value | 0.146-0.353 |

### 3.4. Trajectory Analysis and Structural Stability

To characterize the dynamic behavior of the TRPC6–CAV1 complex and delineate the molecular determinants of their interaction, 300 ns all-atom MD simulations were performed on the modeled assembly. The MD simulations provided detailed insights into the structural dynamics and interaction stability of the TRPC6–CAV1 complex in a cytosolic environment. As shown in (**Fig. 2A**), TRPC6 exhibited pronounced structural flexibility when simulated independently. In contrast, within the TRPC6–CAV1 complex, the backbone RMSD of TRPC6 stabilized between 2.0–2.5 Å, while CAV1 fluctuated moderately within 2.5–3.0 Å (**Fig. 4A**). The RMSD of the overall TRPC6–CAV1 complex remained in the range of 2.5–3.0 Å throughout the trajectory, indicating that both proteins maintain their structural integrity during binding (**Fig. 4A**). This comparable RMSD behavior suggests close structural coupling and dynamic coherence between TRPC6 and CAV1, implying that the two proteins move as an integrated assembly. These results demonstrate that complex formation markedly reduces the intrinsic flexibility of TRPC6, with CAV1 providing stabilizing contacts that suppress large-scale conformational drift and promote a compact, well-organized complex architecture.

**Fig 4.**
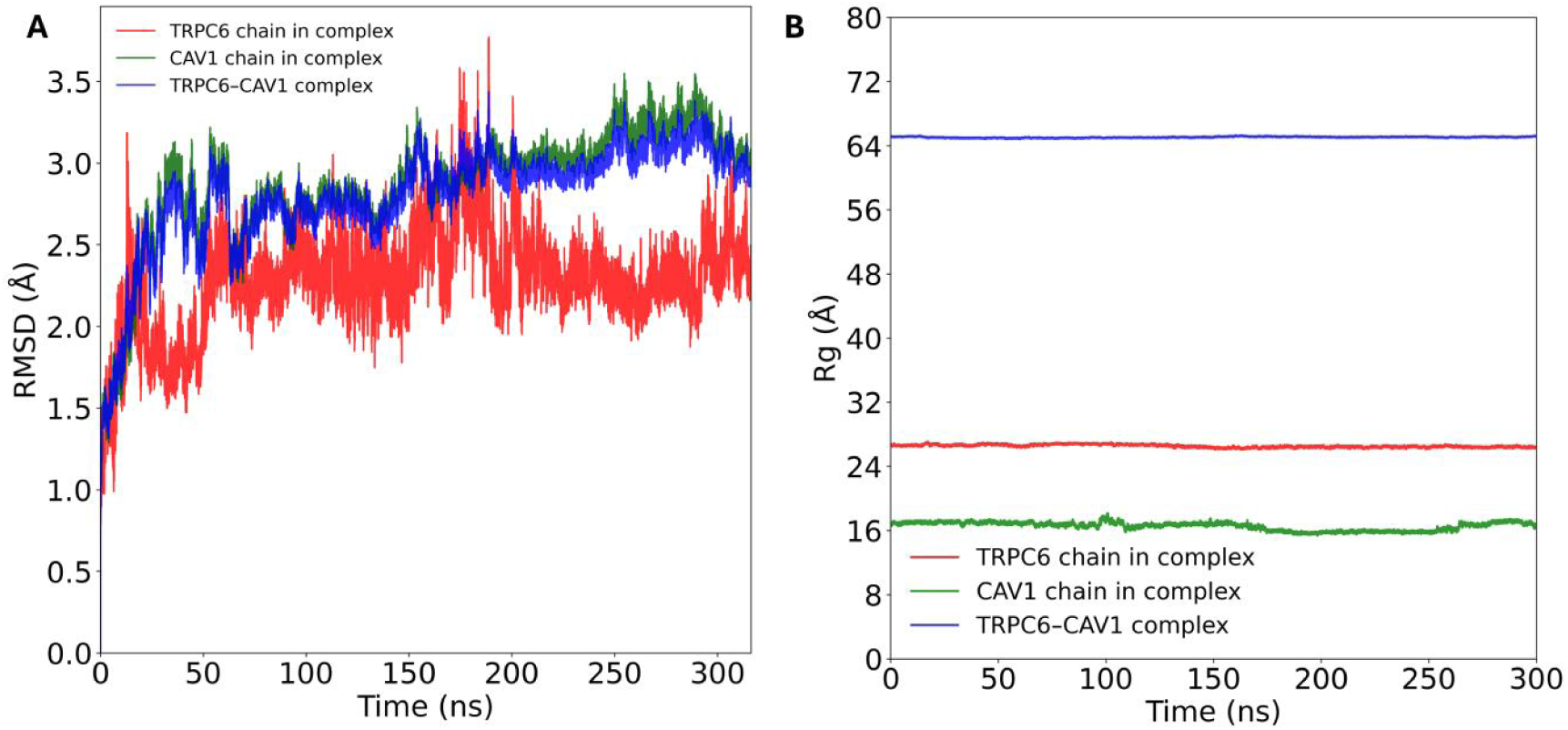
Conformational dynamics of TRPC6 and CAV1. (A) RMSD profiles of TRPC6 alone (red), CAV1 in complex (green), and TRPC6–CAV1 complex (blue) over 300 ns simulation. Complex formation stabilizes TRPC6 compared to the unbound state.

The Rg of TRPC6 remained nearly constant at ∼26.2 Å throughout the 300 ns production phase, indicating that complexation with CAV1 suppresses large-scale fluctuations and stabilizes the TRPC6 architecture. The overall Rg of the TRPC6–CAV1 complex fluctuated around 65 Å across the trajectory, confirming persistent structural integrity without evidence of global collapse or expansion. CAV1 exhibited a comparable stabilization pattern, maintaining an Rg of approximately 17 Å during the simulation, consistent with a stable structural conformation upon binding. The correlated Rg profiles of TRPC6, CAV1, and the complex are depicted in **Fig. 3B**, suggesting synchronized motions and collective stability, supporting the view that the TRPC6– CAV1 assembly functions as a single, dynamically coherent structural unit. Together, these observations highlight the scaffolding role of CAV1 in enhancing TRPC6’s conformational stability and organization, reinforcing its function as a structural regulator rather than an agent of destabilization.

### Solvent-accessible surface area analysis

Solvent-accessible surface area (SASA) analysis (**Fig. 5**) provided dynamic validation of the static interface characterization by quantifying changes in solvent exposure throughout the molecular dynamics trajectory. The total SASA of the TRPC6–CAV1 complex progressively decreased from approximately 135 nm² at t = 0 to ∼130 nm² by 220 ns, corresponding to a ∼5 nm² (3.7%) reduction, before reaching a stable plateau that persisted through 300 ns. This burial of polar and nonpolar surface area reflects progressive interface consolidation upon complex formation and quantitatively corroborates PDBePISA measurements indicating 749.0 Å² and 796.1 Å² of buried surface area for TRPC6 and CAV1, respectively, at the protein–protein contact region. The stable SASA plateau observed over the final ∼80 ns indicates that the interface reached a compact and equilibrated conformation. These time-resolved observations are consistent with PDBePISA-predicted favorable solvation energy gains (ΔⁱG = −2.1 to −5.3 kcal/mol), providing independent dynamic support for the energetic favorability and structural stability of the TRPC6–CAV1 complex in aqueous solution.

**Fig 5.**
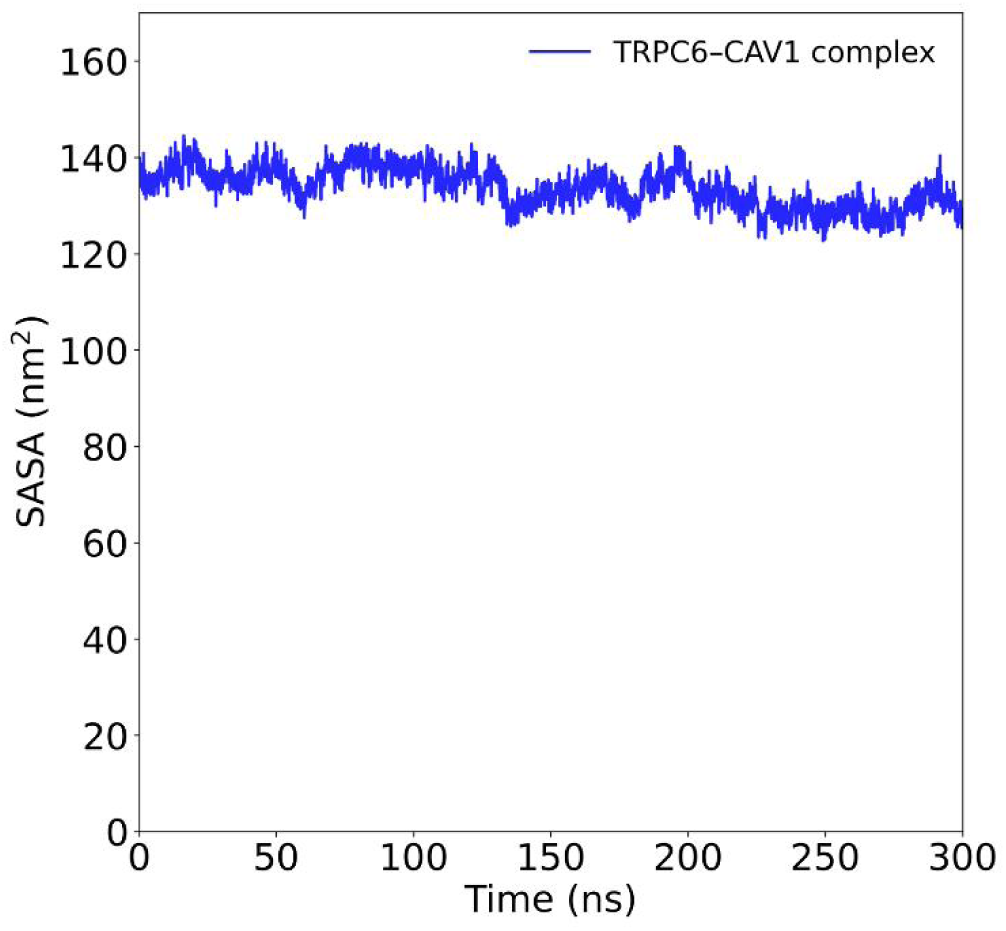
SASA Analysis of TRPC6–CAV1 Complex. Solvent-accessible surface area (SASA) of TRPC6, CAV1, and the complex over the simulation, highlighting interface burial and stabilization of the protein– protein interaction.

### 3.4. Binding Free Energy and Per-Residue Hotspot Analysis

The binding affinity of the TRPC6–CAV1 complex was quantified using MM/PBSA calculations over the equilibrated portion of a 300 ns molecular dynamics trajectory. The total binding free energy (ΔG_binding) was −255.855 ± 1.624 kJ/mol, consistent with a highly stable complex and limited energetic fluctuation across the ensemble. Electrostatic interactions represented the dominant stabilizing contribution (−275.601 ± 1.723 kJ/mol), complemented by a substantial van der Waals term (−34.507 ± 1.071 kJ/mol), whereas polar solvation was unfavorable (+58.657 ± 2.202 kJ/mol) and partially offset by favorable nonpolar (SASA-based) solvation (−4.476 ± 0.015 kJ/mol) in line with typical MM/PBSA behavior for charged protein–protein interfaces. Per-residue MM/PBSA decomposition revealed discrete energetic hotspots at the TRPC6–CAV1 interface (**Fig. 6**).

**Fig. 6.**
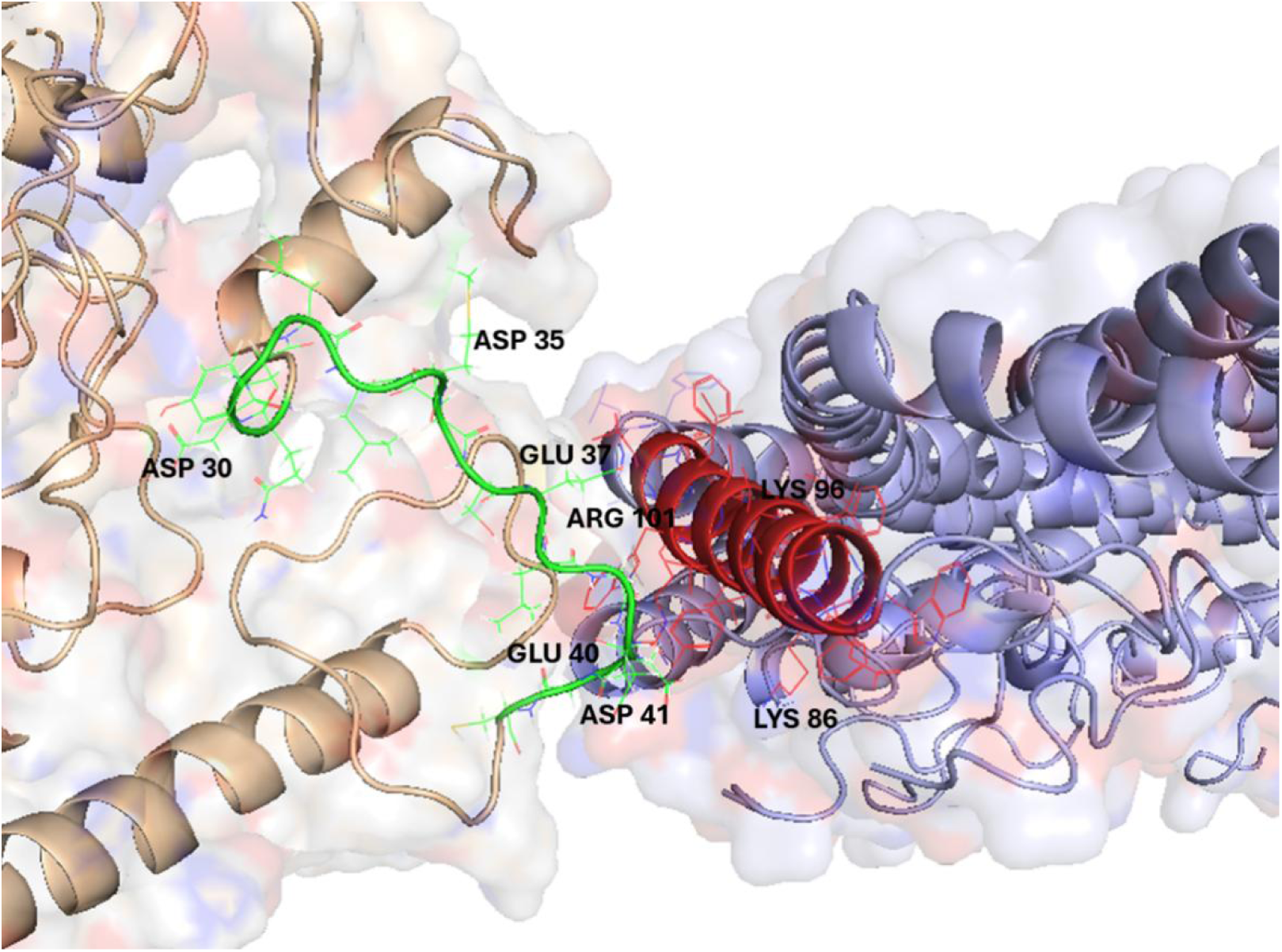
Residues involved in protein–protein interactions. View of the cytosolic interface showing residues participating in TRPC6–CAV1 binding, highlighting key contacts mediating the protein–protein interaction.

In TRPC6, the strongest favorable contributions localized to the cytosolic N-terminal region, forming a contiguous acidic hotspot spanning residues 30–41 (sequence “DYLLMDSELGEGDG”). Within this segment, Asp30 (−18.53 ± 0.13 kJ/mol), Asp35 (−25.22 ± 0.11 kJ/mol), Glu37 (−35.29 ± 0.18 kJ/mol), Glu40 (−38.57 ± 0.30 kJ/mol), and Asp41 (−45.06 ± 0.47 kJ/mol) made the largest stabilizing contributions, defining a negatively charged patch that acts as an electrostatic anchoring surface for CAV1.

A complementary hotspot in CAV1 was identified within residues 85–106 (sequence “WKASFTTFTVTKYWFYRLLSAL”). In this region, Lys86 (−76.87 ± 0.36 kJ/mol), Lys96 (−51.03 ± 0.18 kJ/mol), and Arg101 (−48.66 ± 0.19 kJ/mol) dominated the interaction energetics, underscoring the central role of basic residues in mediating electrostatic attraction to the acidic TRPC6 N-terminus. Additional stabilization arose from hydrophobic and aromatic residues, including Thr90 (−2.98 ± 0.09 kJ/mol), Phe89 (−0.80 ± 0.02 kJ/mol), Val94 (−6.05 ± 0.10 kJ/mol), Phe92 (−0.47 ± 0.02 kJ/mol), Tyr97 (−2.29 ± 0.05 kJ/mol), Trp98 (−5.09 ± 0.14 kJ/mol), and Phe99 (−1.22 ± 0.04 kJ/mol), indicating cooperative contributions from hydrophobic packing and aromatic stacking interactions characteristic of caveolin-mediated recognition. This analysis provides a quantitative estimate of the binding free energy, highlighting the thermodynamic favorability of each residue’s contribution to the complex, as illustrated in **Fig. 7**.

**Fig 7.**
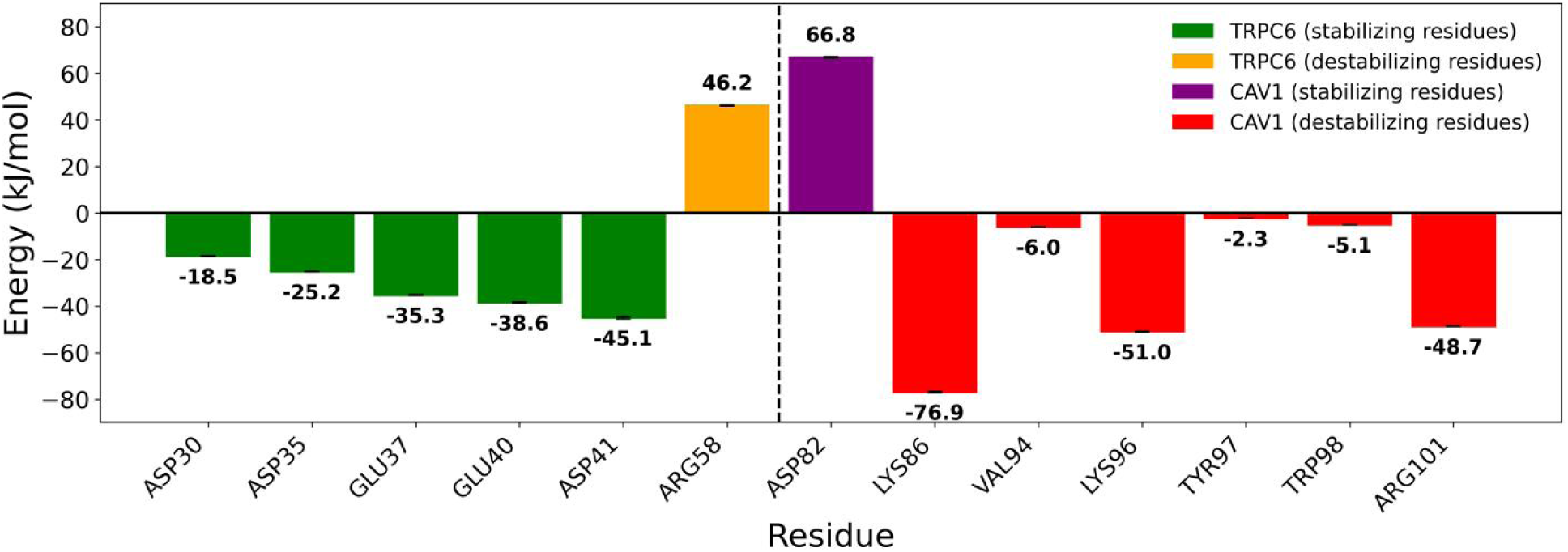
Hot spot per-residue MM-PBSA binding free energy decomposition of the TRPC6–CAV1 complex.

The TRPC6–CAV1 association is mediated exclusively by noncovalent forces, dominated by long-range electrostatic attraction and short-range van der Waals and aromatic stacking, with no evidence of covalent bonding. This interpretation is further supported by the interface and binding analysis of the protein–protein complex in previous section, which confirms that interfacial hydrogen bonds and salt bridges are present but limited in number and do not form a persistent or extensive interaction network. Hydrogen bonds and salt bridges at the interface are transient and continuously exchanged over the simulation timescale, consistent with the dynamic, reversible nature of caveolae-dependent signaling rather than a rigid, permanently locked hydrogen-bond network. The restrained–unrestrained simulation protocol was essential to focus conformational sampling on the biologically relevant cytosolic interface, yielding robust energetic estimates while limiting artifacts such as non-physiological drift or dissociation. These results define a binding interface formed by two complementary energetic hotspots: an acidic N-terminal region in TRPC6 (residues 30–41) and a basic, partially hydrophobic region in CAV1 (residues 85–106). The electrostatically driven and amphipathic nature of this interface suggests a cooperative and dynamic binding mechanism, in which charge complementarity is modulated by transient hydrogen bonding, hydrophobic packing, and aromatic interactions.

## 4. Conclusion

The molecular basis of TRPC6 recruitment and stabilization within caveolar microdomains is fundamental to understanding how spatially restricted calcium signaling is regulated in health and disease. Yet, the previous lack of atomistic information on the TRPC6– CAV1 complex has constrained mechanistic insight into how this interaction is established, maintained, and dynamically tuned. By combining molecular dynamics simulations, MM/PBSA binding free energy calculations, per-residue energy decomposition, and interface analysis, this study provides a detailed atomistic description of the TRPC6–CAV1 interaction. Our results show that TRPC6–CAV1 complex formation is driven primarily by electrostatic interactions, with additional stabilization contributed by van der Waals, hydrophobic, and aromatic contacts that together give rise to a stable yet dynamically adaptable interface. Per-residue hotspot analysis reveals a contiguous acidic patch within the cytosolic N-terminal region of TRPC6 (residues 30–41) that interacts with a complementary basic and partially hydrophobic segment of CAV1 (residues 85–106), defining an amphipathic recognition surface optimized for reversible binding. Interface analysis further indicates that the association is mediated exclusively by noncovalent forces, with hydrogen bonds and salt bridges forming transient, dynamically exchanged contacts rather than a rigid, persistent network—features consistent with the rapid assembly and disassembly of caveolae-associated signaling complexes. Collectively, these findings establish an atomistic framework for TRPC6 regulation by CAV1 within caveolae and highlight the central role of dynamic, reversible protein–protein interactions in caveolin-mediated signaling control. The results of this study provide a rational basis for the development of peptide-based or small-molecule inhibitors targeting the TRPC6–CAV1 interface to selectively disrupt pathological TRPC6 stabilization and calcium signaling, while preserving the essential ion-conducting function of the channel.

## Acknowledgments

The authors would like to thank and acknowledge the lab members of the Universities of Oviedo, and Occupational and Environmental Health. We would also express our gratitude to Jordan Beck and Bernard Monte Pettit for their invaluable comments on modeling TRPC6 complexes.

